# Does late water deficit induce root growth or senescence in wheat?

**DOI:** 10.1101/2023.08.01.551574

**Authors:** Kanwal Shazadi, John T. Christopher, Karine Chenu

## Abstract

Drought frequently limits productivity in rain-fed systems. To investigate water-stress impacts on post-anthesis root development in wheat, three experiments were conducted with two cultivars, Scout and Mace, grown in 1.5m tubes under well-watered conditions or post-anthesis water-stress. Shallow roots of both genotypes appeared to senesce between heading and maturity under well-watered conditions, whereas moderate water stress tended to increase post-heading shallow-root senescence in Mace but stimulated growth in Scout. For deep roots, net growth in biomass was observed for both genotypes under well-watered conditions between heading and maturity, whereas under moderate water stress, only Scout maintained net growth, with net senescence being observed for Mace. Severe water stress resulted in root senescence at all depths for both genotypes. Above ground, Scout retained leaf greenness for only slightly longer than Mace under well-watered conditions. In contrast, under moderate water stress, Mace showed rapid post-anthesis leaf senescence while Scout was affected little if at all. Grain biomass per plant was similar between genotypes in well-watered conditions but more reduced for Mace under moderate stress. Post-anthesis differences in root growth and senescence can strongly influence water use and grain filling in drought-prone environments. Screening for this could assist breeding for drought tolerant varieties.

## 1. Introduction

Wheat cultivation in rain-fed agricultural systems is commonly challenged by water stress during critical stages of plant development (Chenu *et al*., 2013; Farooq *et al*., 2009; Richter and Semenov, 2005). Water stress is a major limiting factor (Araus *et al*., 2002) that commonly limits shoot and root growth (Blum and Sullivan, 1997; Frensch, 1997; Munns, 2002) reducing green leaf area (Christopher *et al*., 2016; Christopher *et al*., 2014; Passioura, 1988) and significant impacting yield. A wide range of physiological and morphological traits have been reported to improve water uptake, water use and water use efficiency in crops like wheat (Christopher *et al*., 2008; Collins *et al*., 2021; Mathew and Shimelis, 2022; Shavrukov *et al*., 2017).

Improved access to water late in the season can be particularly beneficial for wheat crops as under water stress a relatively small amount of subsoil water at this stage can translate to a major yield gain (Kirkegaard *et al*., 2007; Veyradier *et al*., 2013). In wheat, crop simulations have shown that an additional mm of water used after anthesis can lead to an extra 60 kg ha^-1^ of grain yield (Manschadi *et al*., 2010; Manschadi *et al*., 2006). Similar results were observed in field experiments (Kirkegaard *et al*., 2007). While water stress occurring after anthesis has been associated with decreased photosynthetic capacity due to early leaf senescence (Yang *et al*., 2001), genotypes with the ability to retain green leaf area during the grain filling period, i.e. with a stay-green phenotype, tend to yield more than others in this type of environment (Christopher *et al*., 2016; Christopher *et al*., 2014; Kumar *et al*., 2010). Comparing two cultivars with contrasting stay-green pattern, Christopher et al. (2008) found that those genotypes also differed in terms of root architecture, particularly at depth. In large soil-filled chambers, the drought tolerant line SeriM82 compared with the Australian cultivar Hartog was found to have a narrower root architecture with a greater proportion of roots at depth, which resulted in greater soil water extraction from deep in the profile (Manschadi *et al*., 2006). However, although higher root biomass and root distribution deeper in the profile has commonly been associated with better adaptation to water stress, little is known about the dynamics of root development late in the crop cycle and under water stress (Palta *et al*., 2011).

A well-adapted root architecture can improve water availability for the crop by increasing the amount of water that can be extracted from the soil. The dynamic of root development can also be important to allow water uptake at critical stages (e.g. Veyradier *et al*., 2013). Beneficial root traits vary depending on the target population of environments. It has been suggested that in environments with shallow soils and frequent low-intensity rainfall, developing a dense root system in shallow soil layers may be advantageous to crops (Gregory *et al*., 1978; López-Castañeda and Richards, 1994). By contrast, in environments where crops rely heavily on water stored deep in the soil such as in north-eastern Australia (Chenu *et al*., 2011 and 2013), a deep root system may allow crops to perform better under late water stress. In such environments, root traits reported to be beneficial include higher root-length density at depth (Lopes and Reynolds, 2010) and more uniform root distribution at depth (Manschadi *et al*., 2006) to allow crops to extract additional soil water at depth (Asseng and Turner, 2007; Lilley and Kirkegaard, 2011; Ober *et al*., 2014), keep the canopy functional (Christopher *et al*., 2008) and increase yield (Dodd *et al*., 2011). The dynamic of growth and senescence of those traits is also important, as this influences the dynamic of water uptake.

The aim of this study was to characterise post-heading root development over time for two contrasting wheat cultivars in well-watered and water-stressed conditions. The root system of Scout and Mace, which are known to differ in seedling root traits, were examined at key stages from heading to maturity in well-watered conditions and a range of post-anthesis water stress treatments. A system of 1.5 m polyvinyl chloride (PVC) tubes was used to investigate phenotypic differences in both above- and below-ground response to water stress up to physiological maturity.

## 2. Materials and Methods

### 2.1 Growing conditions and experimental design

Three experiments were conducted with two wheat cultivars (Mace and Scout) grown in 1.5-m long polyvinyl chloride (PVC) tubes of 90 mm diameter under different soil water conditions. Tubes were placed in an outdoor open area in Toowoomba, Queensland, Australia (27.5598°S, 151.9507°E, 691 meters a.s.l.).

Mace and Scout are widely cultivated in western and southern cropping regions of Australia, respectively and differ in root architecture at the seedling stage, as Mace has wide root angle while Scout has narrow root angle (Richard *et al*., 2018). Three seeds of each genotype were placed in each tube at a depth of 2 cm. Following emergence, plants were thinned to one seedling per tube.

Seeds were sown in a packed soil consisting of a 50:50 mixture of a red alluvial soil from Redlands (−27.53°S, 153.25°E, ∼20 m a.s.l.) and black vertosol soil from Kingsthorpe (27.51°S, 152.10°E, ∼480 m a.s.l.). To ensure non-limiting nutrient supply, 2 gm L^-1^ of Osmocote fertilizer containing trace elements (N 15.3% P 1.96%, K 12.6%) was added to the soil mix. The soil was watered to field capacity at sowing.

Experimental treatments were denominated by the experiment number (E1, E2, E3) followed by “WW” for well-watered, “MDE” for moderate drought early in grain filling, “MDM” for moderate drought mid-grain filling or “SD” for severe drought with water being withheld from head emergence to maturity (Table 1). Growth stages of individual replicate plants were monitored, and watering withheld between the required developmental periods (Table 1).

**Table 1.** Characteristics for experiments (Exp) and treatments (Tmt) including the Zadoks developmental stages between which irrigation was withheld, number of days from sowing to anthesis (Anthesis), from sowing to maturity (Maturity), and from anthesis to maturity (Grain filling period), green leaf area of the plant at heading (Z50) and grain yield per plant at maturity (Z92) for Mace and Scout. Average and standard errors were presented for days to anthesis and maturity as well leaf area and yield (n=8). Means followed by different superscript letters are significantly different between genotypes within each treatment (*P*<0.05), “ns” indicates that the difference between genotypes is not significant (*P*>0.05). Each experiment was analyzed separately.

| Experiment | Sowing | Tmt* | Water withholding period** | Genotype | Anthesis<br>(Z65)<br>(days) | Maturity<br>(Z92)<br>(days) | Leaf area<br>(Z50)<br>(cm<sup>2</sup> plt<sup>-1</sup>) | Yield<br>(Z92)<br>(g plt<sup>-1</sup>) |
| --- | --- | --- | --- | --- | --- | --- | --- | --- |
| Experiment 1 | 4/08/2020 | E1- WW | none | Mace | 77±0.9<sup>ns</sup> | 129±0.8<sup>ns</sup> | 684±55<sup>b</sup> | 16±1.5<sup>ns</sup> |
|  |  |  |  | Scout | 83±0.8<sup>ns</sup> | 131±0.9<sup>ns</sup> | 834±78<sup>a</sup> | 14±1.5<sup>ns</sup> |
|  | 4/08/2020 | E1-MDE | Z60-71 | Mace | 75±0.9<sup>b</sup> | 117±0.8<sup>b</sup> | NA | 7±1.6<sup>b</sup> |
|  |  |  |  | Scout | 80±0.8<sup>a</sup> | 133±0.9<sup>a</sup> |  | 16±1.8<sup>a</sup> |
|  | 4/08/2020 | E1-MDM | Z71-81 | Mace | 78±0.9<sup>b</sup> | 117±0.8<sup>b</sup> | NA | 8±1.8<sup>b</sup> |
|  |  |  |  | Scout | 85±0.8<sup>a</sup> | 131±0.9<sup>a</sup> |  | 12±1.5<sup>a</sup> |
| Experiment 2 | 24/08/2021 | E2-WW | none | Mace | 68±1.9<sup>ns</sup> | 106±1.9<sup>ns</sup> | NA | 12±0.4<sup>ns</sup> |
|  |  |  |  | Scout | 74±2.2<sup>ns</sup> | 110 ±2.2<sup>ns</sup> |  | 13±0.5<sup>ns</sup> |
|  | 24/08/2021 | E2-MDE | Z60-71 | Mace | 65±1.9<sup>b</sup> | 96±1.9<sup>b</sup> | NA | 8±0.4<sup>ns</sup> |
|  |  |  |  | Scout | 75±2.2<sup>a</sup> | 119±1.9<sup>a</sup> |  | 14±0.4<sup>ns</sup> |
| Experiment 3 | 4/07/2019 | E3-SD | Z50-92 | Mace | 79±0.8<sup>ns</sup> | 115±0.78<sup>ns</sup> | 395±48<sup>ns</sup> | 2.5±0.19<sup>ns</sup> |
|  |  |  |  | Scout | 86±0.8<sup>ns</sup> | 120±0.78<sup>ns</sup> | 573±48<sup>ns</sup> | 3±0.19<sup>ns</sup> |

\*Experimental treatments (Tmt) were denominated by the experiment number followed by “WW” for well-watered, “MDE” for moderate drought early in grain filling, “MDM” for moderate drought mid-grain filling or “SD” for severe drought during grain filling.
\*\*Period of withholding watering are indicated by the number of the Zadoks growth stage from when watering was discontinued followed by the stage where watering was recommenced.

**Table 2.** Environmental conditions in the three experiments (Exp.), including average temperature (Avg. Temp.), average daily maximum temperature (Tmax), average daily minimum temperature (Tmin), average evaporation (Avg. Evap.), average daily radiation (Avg. Radn) and cumulated radiation (Cum. Radn) from sowing to maturity of the last maturing plant.

| Exp. | Avg. Temp.<br>(°C) | Tmax<br>(°C) | Tmin<br>(°C) | Avg. Evap.<br>(mm) | Avg. Radn<br>(W m<sup>-2</sup>) | Cum. Radn<br>(W m<sup>-2</sup>) |
| --- | --- | --- | --- | --- | --- | --- |
| E1 | 19 | 25 | 13 | 5.1 | 19 | 20.1 |
| E2 | 18.5 | 24 | 13.2 | 4.5 | 18.5 | 19.4 |
| E3 | 17 | 22.5 | 10 | 4.6 | 17 | 18.4 |

In the first experiment (E1), three irrigation treatments were applied; (i) well-watered conditions during the whole crop cycle (E1-WW); (ii) well-watered conditions followed by a water deficit applied by withholding irrigation between early anthesis (Zadoks decimal growth stage 61; Z61) (Zadoks *et al*., 1974) and early grain filling (Z71) (E1-MDE), and (iii) a water deficit imposed by withholding irrigation from early grain filling (Z71) to mid-grain filling (Z81) (E1-MDM). In this experiment, plants were harvested at heading (Z50), anthesis (Z65) and maturity (Z92). In a second experiment (E2), two irrigation treatments were applied; (i) well-watered conditions during the whole crop cycle (E2-WW); (ii) well-watered conditions followed by a water deficit applied by withholding irrigation between early anthesis (Z61) and early grain filling (Z71) (E2-MDE). In this experiment, plants were harvested at mid-grain filling (Z75) and maturity (Z92). In a third experiment (E3), water was withheld for the whole period from head emergence (Z50) to maturity (Z92) (E3-SD).

For all experiments, tubes were watered weekly up to saturation, and watering was stopped for each individual tube one day prior to the target growth stage. For each experiment, a randomized complete block design was used with eight replicates per cultivar for each treatment, a replication being a single plant in a tube.

### 2.2. Plant measurements

Phenological development was monitored regularly by recording the growth stage using the Zadoks growth scale throughout the experiments (Zadoks *et al*., 1974). The greenness of the center of the flag leaf of the main stem was measured at Z50, Z65, Z75 and Z81 using a Minolta SPAD 502 meter (Konica Minolta, Tokyo).

For each harvest, the shoots were excised at the crown. To maintain the root distribution, roots were washed and recovered on a nail board, with nails spaced every 20 mm. Root sections were excised at 10-cm intervals for measurement of dry root biomass. The root biomass was measured following drying for 72h at 70°C. Total root biomass was calculated as the sum of dry weights for all 10 cm samples for each core. Average root diameter and root length were measured for a subset of soil layers (0-10 cm, 10-20 cm, and alternate 10-cm depth intervals there after (i.e., 30-40, 50-60, 70-80, 90-100, 110-120, and 130-140 cm) using WinRhizo Regular 2019. For each 10-cm depth, average root length density was calculated by dividing the total root length in a segment by the corresponding soil volume. To measure the differences in the partitioning of biomass between shallow, mid and deep roots, root fractions from 0-50 cm were summed to represent shallow roots, 50-100 cm to represent mid roots, and 100-150 cm to represent deep roots. Similarly, average root diameter and root length density were estimated for shallow, mid and deep roots by averaging values from studied depths (0-10, 10-20 and 30-40 cm for shallow roots; 50-60, 70-80 cm for mid roots; and 90-100, 110-120 and 130-140 cm for deep roots). The root: shoot ratio was computed by dividing total dry root biomass by the total dry shoot biomass. The total plant biomass was computed by combining the total dry shoot and dry root biomass. For harvests at heading (Z50) and anthesis (Z65), leaf blades were separated for measurement of green leaf area using a leaf area meter (LI-3000, Li-COR Bioscience, Lincoln, NE, USA). Yield was recorded at maturity as the total grain biomass (air dried) per plant.

### 2.3 Statistical analysis

Within each experiment, an analysis of variance (Figueroa-Bustos *et al*., 2020) was performed between genotypes, treatments and stages for total plant biomass, dry shoot biomass, total dry root biomass, average root length density, average root diameter, dry root biomass at depths, average root length density at depths and average root diameter at depths using the R platform (v3.2.5; R Core Team, 2019). A Student–Newman–Keuls (SNK) test was used to compare means for genotypes and treatments, with a significance level of 0.05.

## 3. Results

### 3.1 Water stress reduced the duration of the plant growth cycle in Mace but not in Scout

Under well-watered conditions, the plant growth duration from sowing to anthesis and to maturity was slightly shorter for Mace than for Scout, but differences were not significant (Table 1).

Application of a post-anthesis moderate water stress shortened the duration to maturity in Mace but not Scout. While under well-watered conditions the difference in time to maturity between genotypes was relatively small at 2 or 4 days in Experiments 1 or 2, respectively, this difference ranged from 14 to 23 days under moderate stress (Table 1).

For the severe stress (E3-SD), the 5d difference between genotypes was not significant and relatively small compared to moderately water-stressed plants in the other two experiments.

### 3.2 Under well-watered conditions, dry root biomass at depths increased post anthesis for Scout but not for Mace

Under well-watered conditions, whole-plant dry biomass increased from anthesis to maturity for both Scout and Mace (Fig. 1A), with root biomass being only 2.5-10 % of the shoot biomass (Fig. 1B-D). Mace tended to have a smaller total plant biomass than Scout both at anthesis and maturity (Fig. 1A) mainly due to a smaller shoot system. Mace also tended to have lesser root biomass than Scout, especially at maturity (Z92; Fig. 1C).

**Fig. 1.**
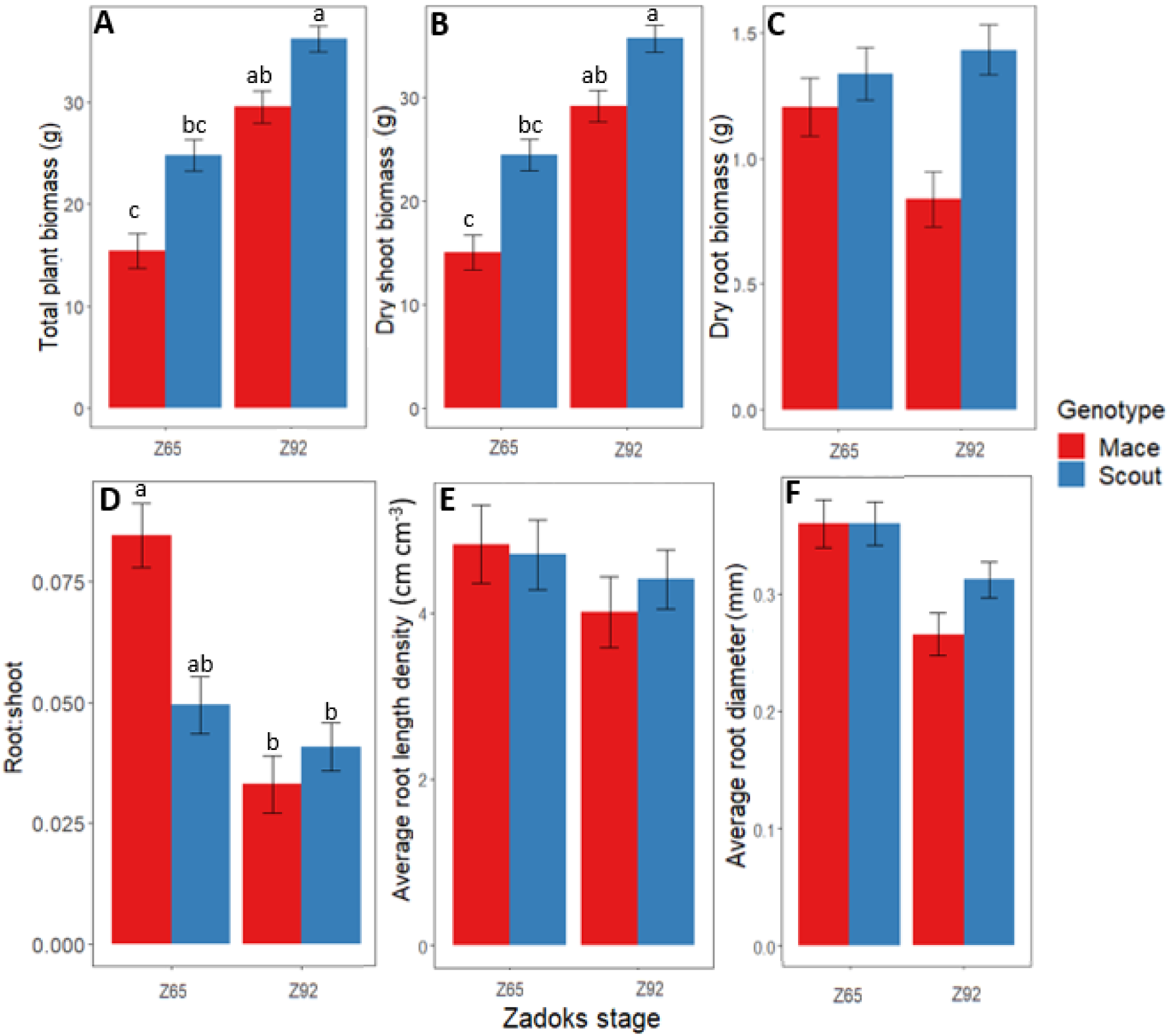
Changes between anthesis (Z65) and maturity (Z92) under well-watered conditions (E1-WW) in (A) whole-plant dry biomass, (B) shoot dry biomass, (C) root dry biomass, (D) root: shoot dry mass ratio, (E) average root length density, and (F) average root diameter for Mace (red bars) and Scout (blue bars). Different letters indicate mean values that are significantly different at P < 0.05. Error bars represent the standard error of the mean (n=8).

Significant post-anthesis root growth occurred for roots deeper than 50 cm in Scout under well-watered conditions (Fig. 2A, D), with a net increase in root biomass between anthesis and maturity (Fig. 2A, D). In contrast, for Mace, root biomass changed little from anthesis and maturity at all depths deeper than 50 cm, while some reduction in root biomass was observed for the shallow soil layers (Fig. 2A, D).

**Fig. 2.**
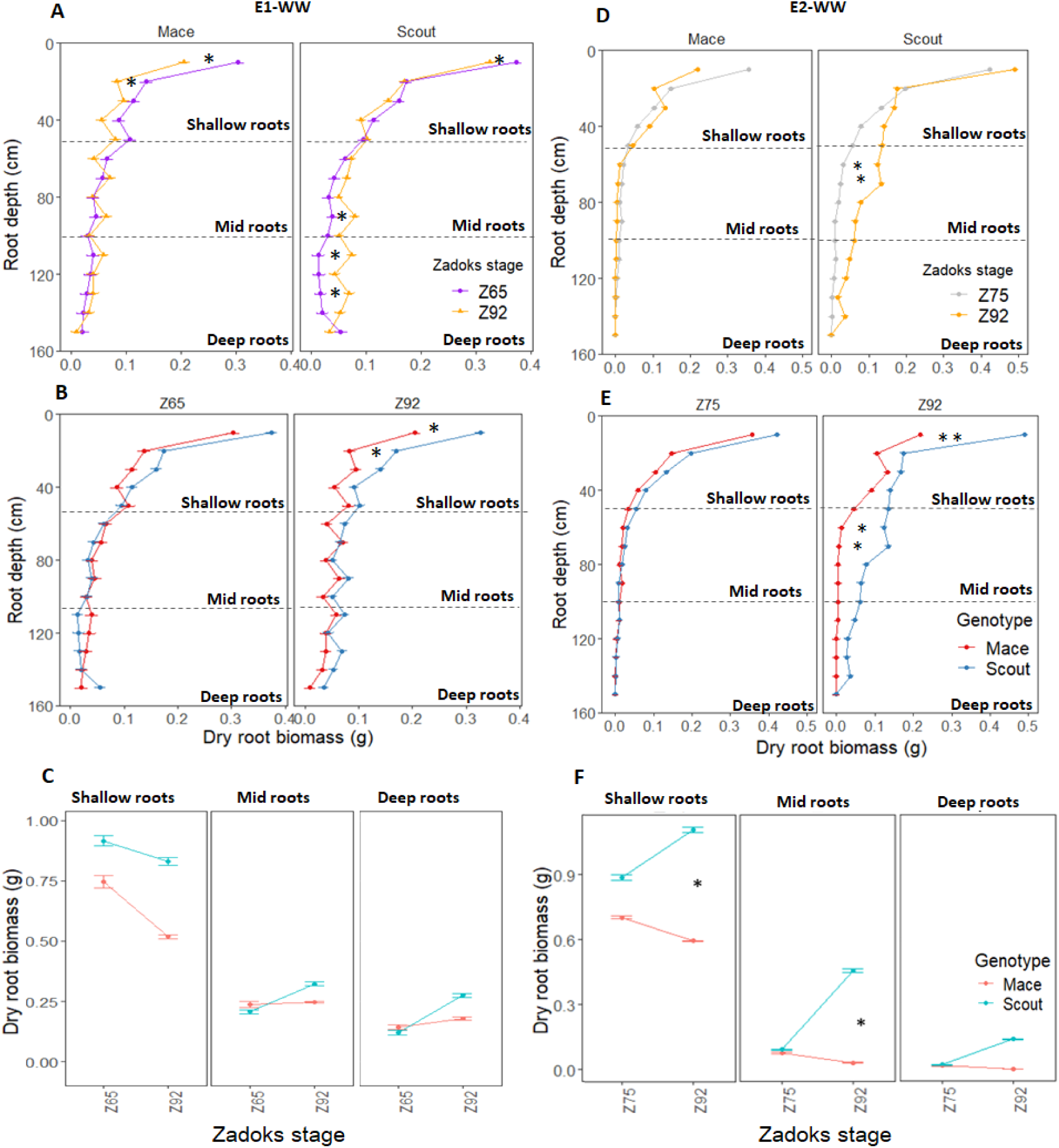
Dry root biomass at different depths (0 to 150 cm) for Mace and Scout plants grown under well-watered conditions in E1-WW (A, B, C) and E2-WW (D, E, F) and harvested at different stages, including anthesis (Z65), mid-grain filling (Z75) and maturity (Z92). In (A-B, D-E), the horizontal dashed lines represent partitions between shallow (0 to 50 cm), mid (50 to 100 cm), and deep (100 to 150 cm) root layers. Error bars represent the standard error of the mean (n=8). Significant differences between means for root layers are indicated by asterisks for P<0.05 * and P<0.01**.

Overall, the post-anthesis increase in shoot biomass with little or no increase in root biomass led to a decrease in the root: shoot ratio from anthesis to maturity for both genotypes (Fig. 1D).

Post-anthesis root length density tended to decrease for shallow roots (< 50 cm) in both genotypes, while it tended to increase in deep roots (> 50 cm) in Scout (Fig. 3A, C). For both genotypes, average root diameter tended to decrease after anthesis at all depths under well-watered conditions (Fig. 3B, D).

**Fig. 3.**
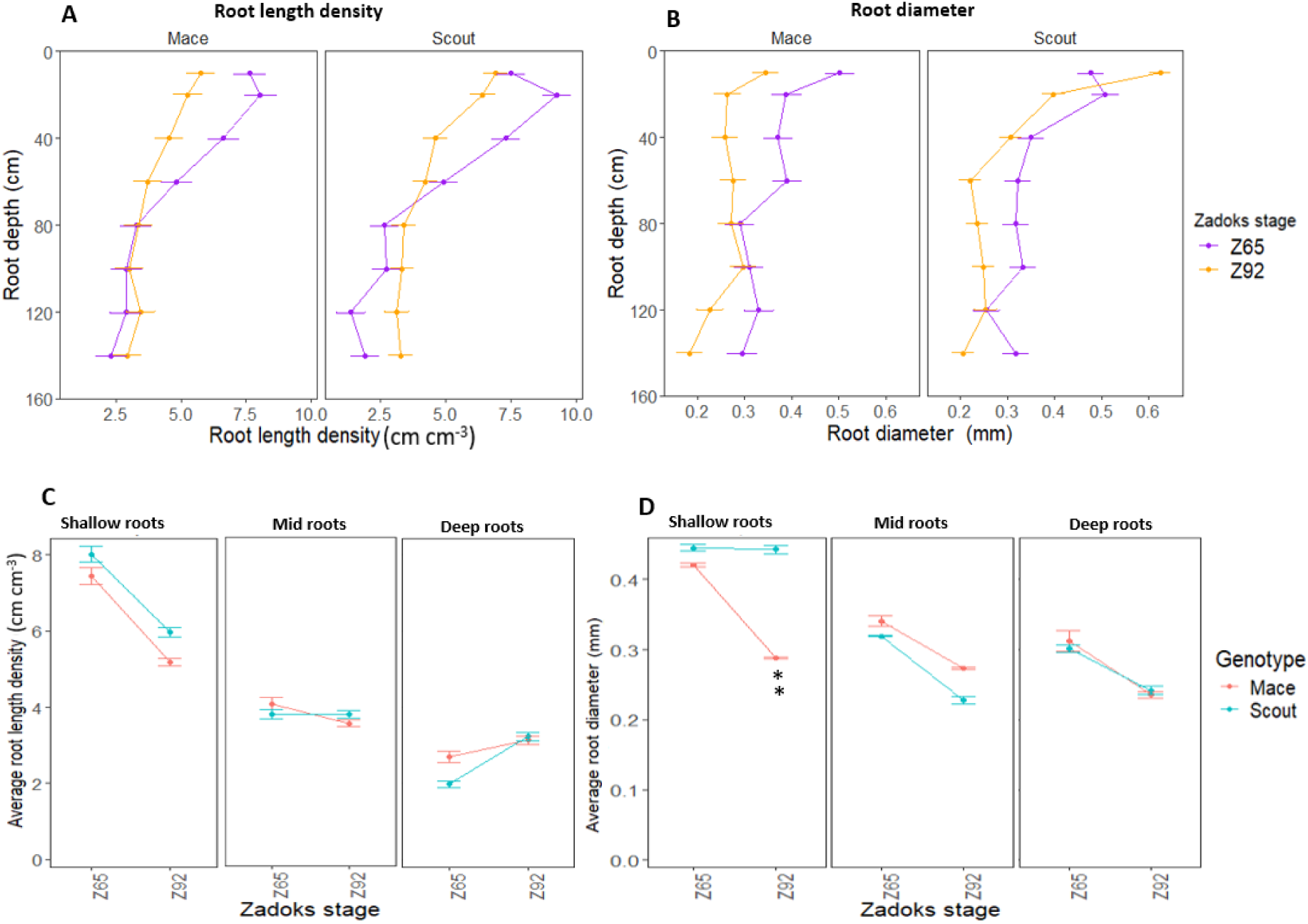
Average root length density (A, C) and root diameter (D, E) of Mace and Scout at anthesis (Z65) and maturity (Z92) under well-watered conditions (E1-WW) at different 1 0-cm depth intervals (0-10, 20-30, 50-60, …, 130-140 cm) (A, B), and for corresponding shallow (0 to 50 cm), mid (50 to 100 cm) and deep (100 to 150 cm) roots (C, D). Error bars represent the standard error of the mean (n=8). **, significant genotypic difference at P<0.01.

### 3.3 Moderate water stress reduced both shoot and root biomass in Mace, while Scout was more tolerant

Total plant biomass at maturity in well-watered conditions was not significantly different between Scout and Mace although Mace tended to have a lower whole-plant dry biomass (Fig. 4A). In contrast, significant differences were observed between genotypes in moderately water-stressed treatments (Fig. 4A). Moderate water-stress treatments during the early or mid-grain filling period significantly reduced whole-plant biomass at maturity in Mace, with a reduction by 48.8%, 37.6% and 32.78% for E1-MDE, E1-MDM, and E2-MDE, respectively, compared to their respective well-watered controls (E1-WW and E2-WW; Fig. 4A). By contrast, whole-plant biomass of Scout was little affected by moderate water stress and remained similar to that observed in well-watered conditions. This distinction between genotypes was lost in the severe water stress treatment (E3-SD) in experiment E3, which substantially reduced the total plant biomass of both genotypes to similarly low values.

**Fig. 4.**
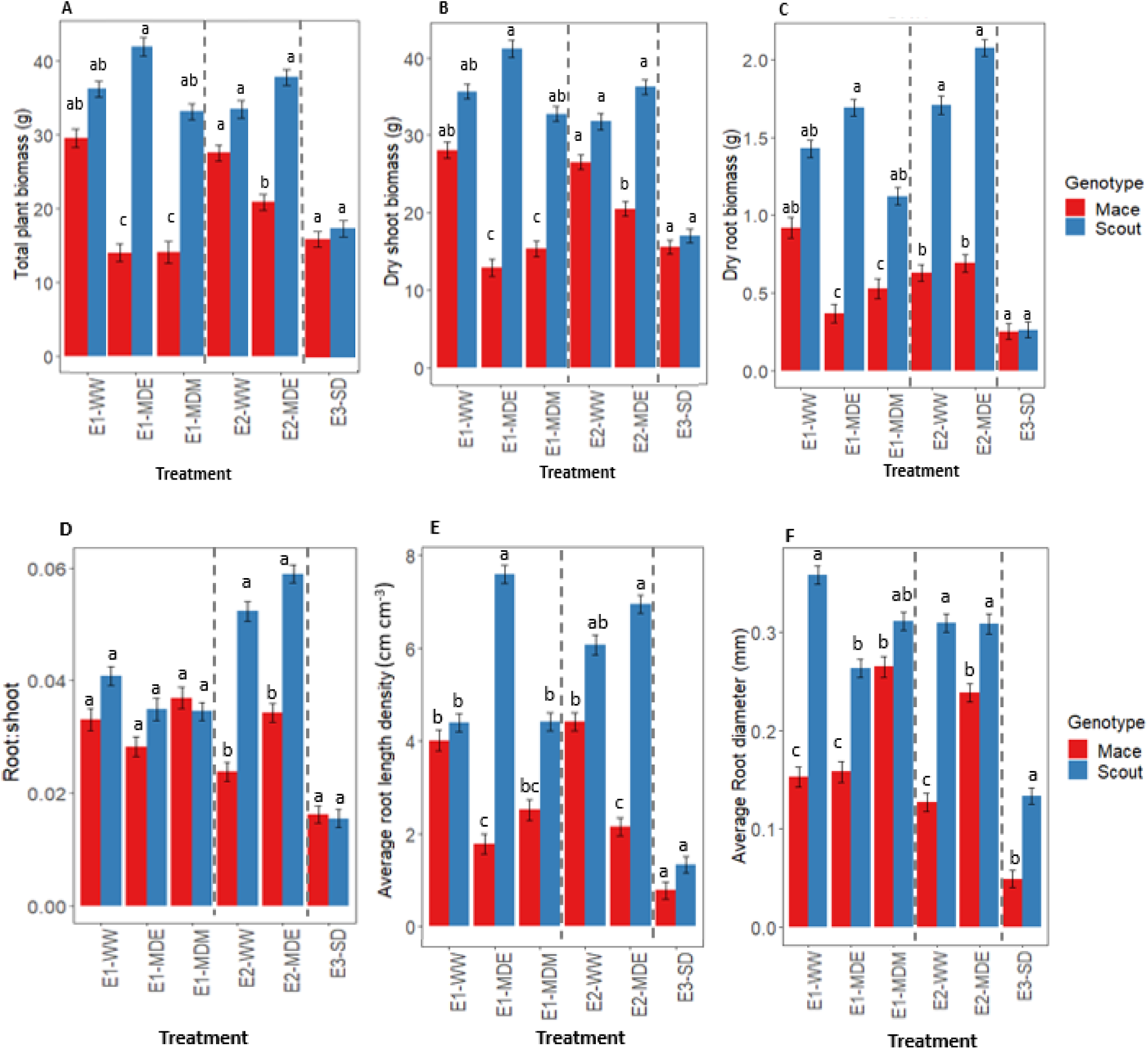
Response to different soil water status at maturity for (A) whole-plant dry biomass, (B) shoot dry biomass, (C) root dry biomass, (D) root: shoot dry mass ratio, (E) average root length density, and (F) average root diameter in Mace (red bars) and Scout (blue bars). In each panel, dashed lines separate the data from the three experiments. Analysis of variance was performed separately for each experiment. Different letters indicate mean values that are significantly different at P < 0.05 within each experiment. Error bars represent the standard error of the mean (n=8).

A similar trend was observed for dry shoot biomass with greater difference between the genotypes for moderate water stress treatments (E1-MDE, E1-MDM and E2-MDE) compared to either the well-watered or severe water stressed treatments (Fig. 4B).

Root biomass and root length density also followed an overall similar trend (Fig. 4C, E). Average root diameter was greater in Scout than in Mace in most treatments, including under well-watered conditions. By contrast, a moderate water stress had either no effect or increased the average root diameter in Mace, while it decreased average root diameter in Scout in experiment E1 (Fig. 4F).

The root: shoot ratio at maturity did not vary significantly between treatments or between genotypes in experiment E1 (Fig. 4D). However, in experiment E2, significant differences were observed between genotypes both in E2-WW and E2-MDE, with Scout having a greater root: shoot ratio than Mace (Fig. 4D). For the severe water-stress treatment of experiment E3 (E3-SD), no significant difference was observed between Scout and Mace for root: shoot ratio. The mean values for root: shoot ratio, although they cannot be formally compared between experiments, were much lower in E3 than in any of the treatments in E1 and E2.

### 3.4 Moderate water stress induced root senescence at all depths in Mace, and in shallow roots in Scout

For Mace, no significant difference was observed in root biomass between water treatments at shallow depths (0-50 cm), but moderate water stress (E1-MDE, E1-MDM and E2-MDE) caused root senescence for lower depths (100-150 cm) compared to well water treatments (E1-WW and E2-WW; Fig. 5, Supplementary Fig. S1). By contrast in Scout, no substantial difference in root biomass was observed for deep roots between well-watered (E1-WW and E2-WW) and moderate water stress treatments. However, shallow roots of Scout after a moderate stress during mid-grain filling (E1-MDM) or a severe stress (E3-SD) had less biomass than well-watered plants at maturity, indicating root senescence after flowering. By contrast, for an earlier moderate stress during early grain filling (E1-MDE and E2-MDE), Scout shallow roots tended to grow more than under the other treatments, including well-watered conditions (E1-WW and E2-WW). In the severely water stressed treatment (E3-SD), dry root biomass for shallow and deep roots was similarly severely reduced for both genotypes at maturity.

**Fig. 5.**
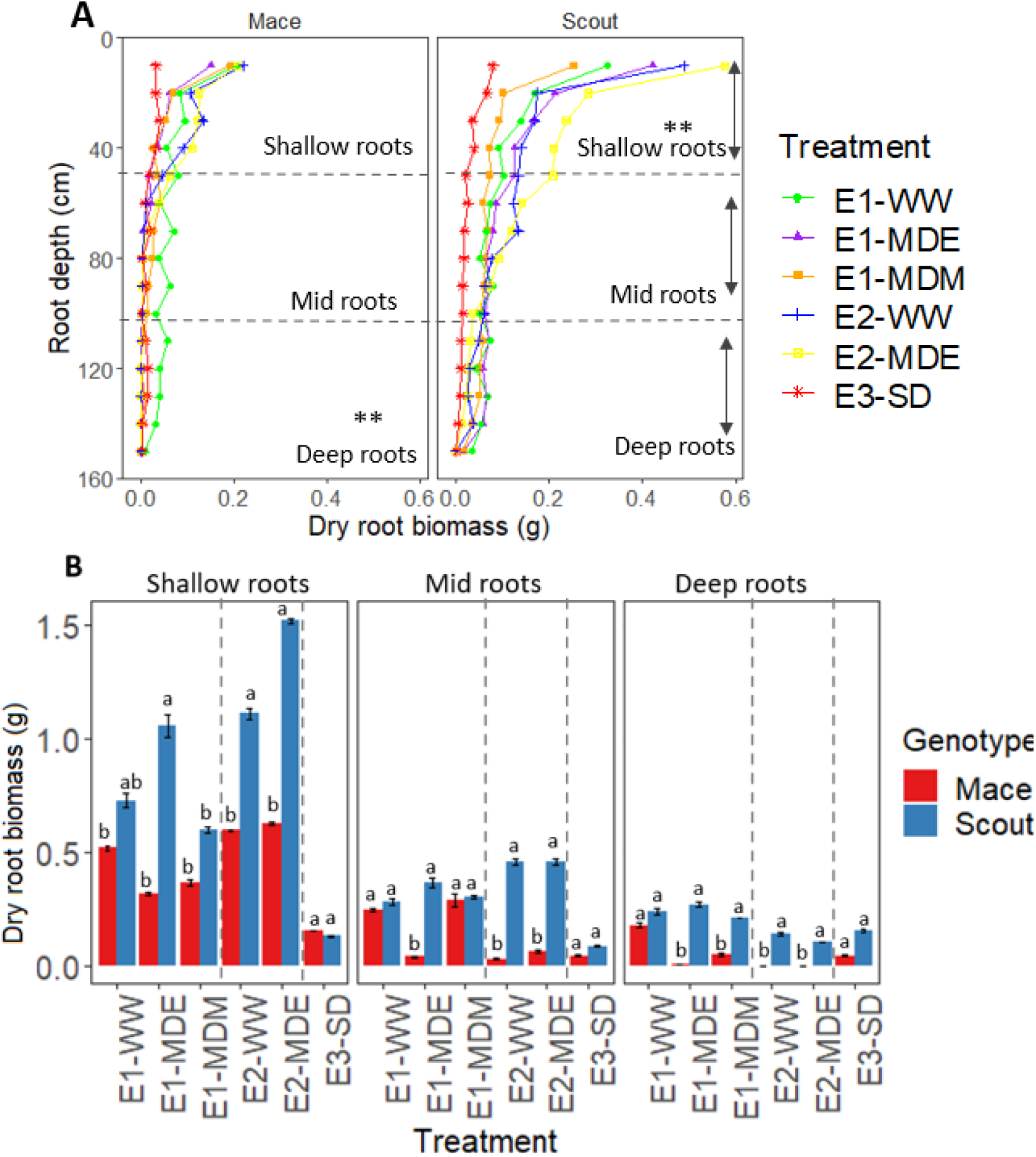
Distribution of dry root biomass for different soil water treatments at maturity for (A) 10-cm increment depths (0 to 150 cm) and (B) shallow (0 to 50 cm), mid (50 to 100 cm), and deep (100 to 150 cm) roots in Mace and Scout. In (A), asterisks indicate differences between treatments for dry biomass of shallow, mid or deep roots (P<0.001). In (B), the dotted lines separate the three independently analysed experiments; means that are significantly different (P<0.05) within each experiment are indicated by different letters above the bars. Error bars represent the standard error of the mean (n=8).

Root length density and average root diameter for shallow and deep layers had a similar trend to that of root biomass under all treatments for both Scout and Mace (Fig. 6, Supplementary Figs. S2, S3).

**Fig. 6.**
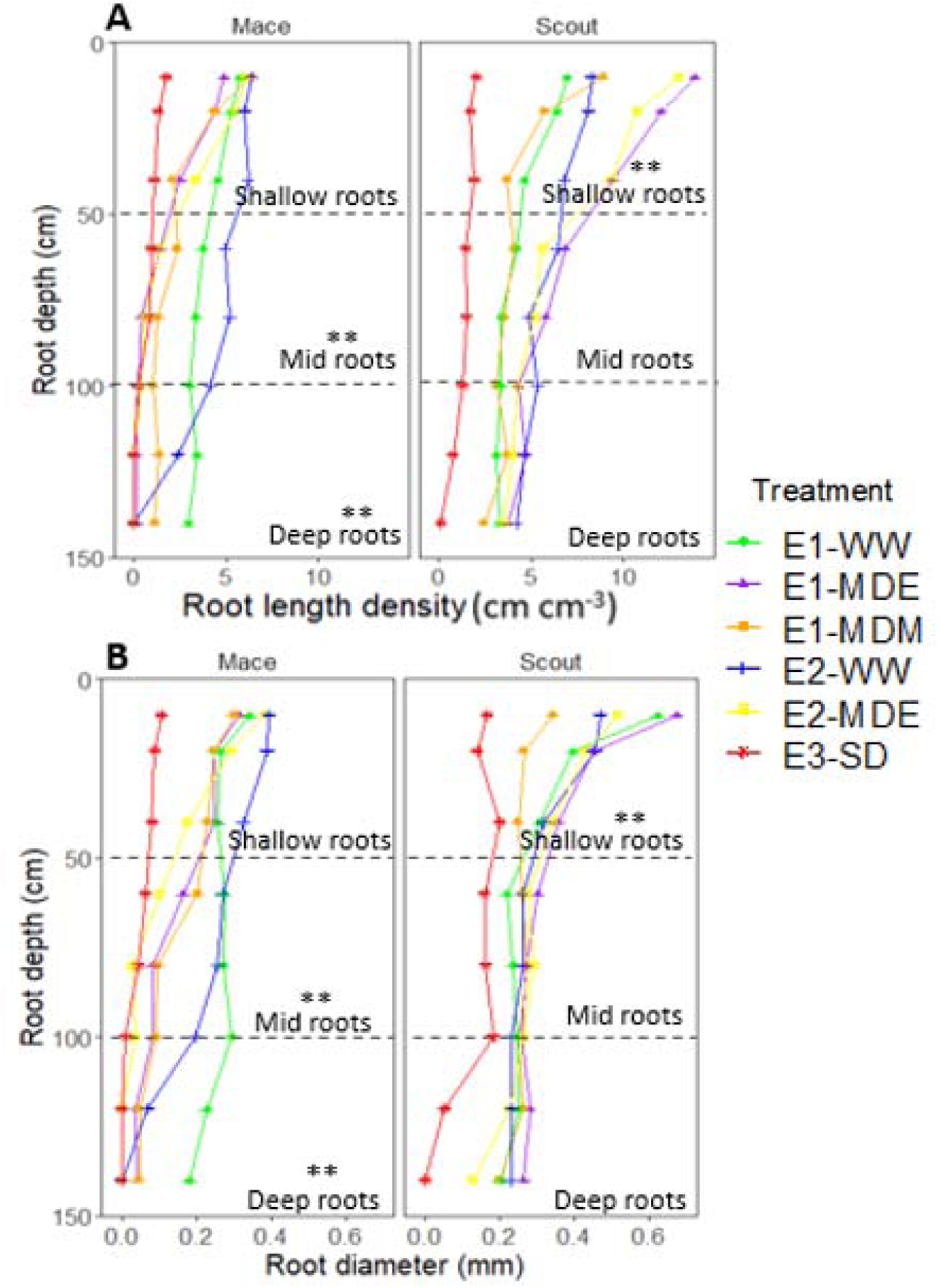
Root length density (A) and root diameter (B) of Mace and Scout for different depths (0 to 150 cm) at maturity for different water status treatments. The dashed lines represent partitions between shallow (0 to 50 cm), mid (50 to 100 cm), and deep (100 to 150 cm) roots. Asterisks indicate differences between treatments for shallow, mid or deep roots (P<0.01).

To investigate dynamic changes in root development, variations in dry root biomass distribution between heading (Z50) and maturity (Z92) were estimated by subtracting the root biomass at heading from the root biomass at maturity (Fig. 7A, Supplementary Fig. S4), completing comparison previously done between anthesis and maturity (e.g., Figs 1, 2A-C, 3). For Mace, all water-stress treatments tended to cause net shallow and deep root senescence between heading and maturity, while in well-watered conditions, net senescence was only observed in shallow roots and to a lesser extent than for stressed conditions (Fig. 7A). In contrast for Scout, deep root biomass increased between heading and maturity in all treatments, except for the severe water-stress treatment (E3-SD) where net senescence was observed. For Scout shallow roots, senescence was observed for all treatments except E1-MDE. In other words, deep roots for Scout grew between heading and maturity in all studied conditions, except the severe stress E3-SD which induced a net root senescence in all soil layers.

**Fig. 7.**
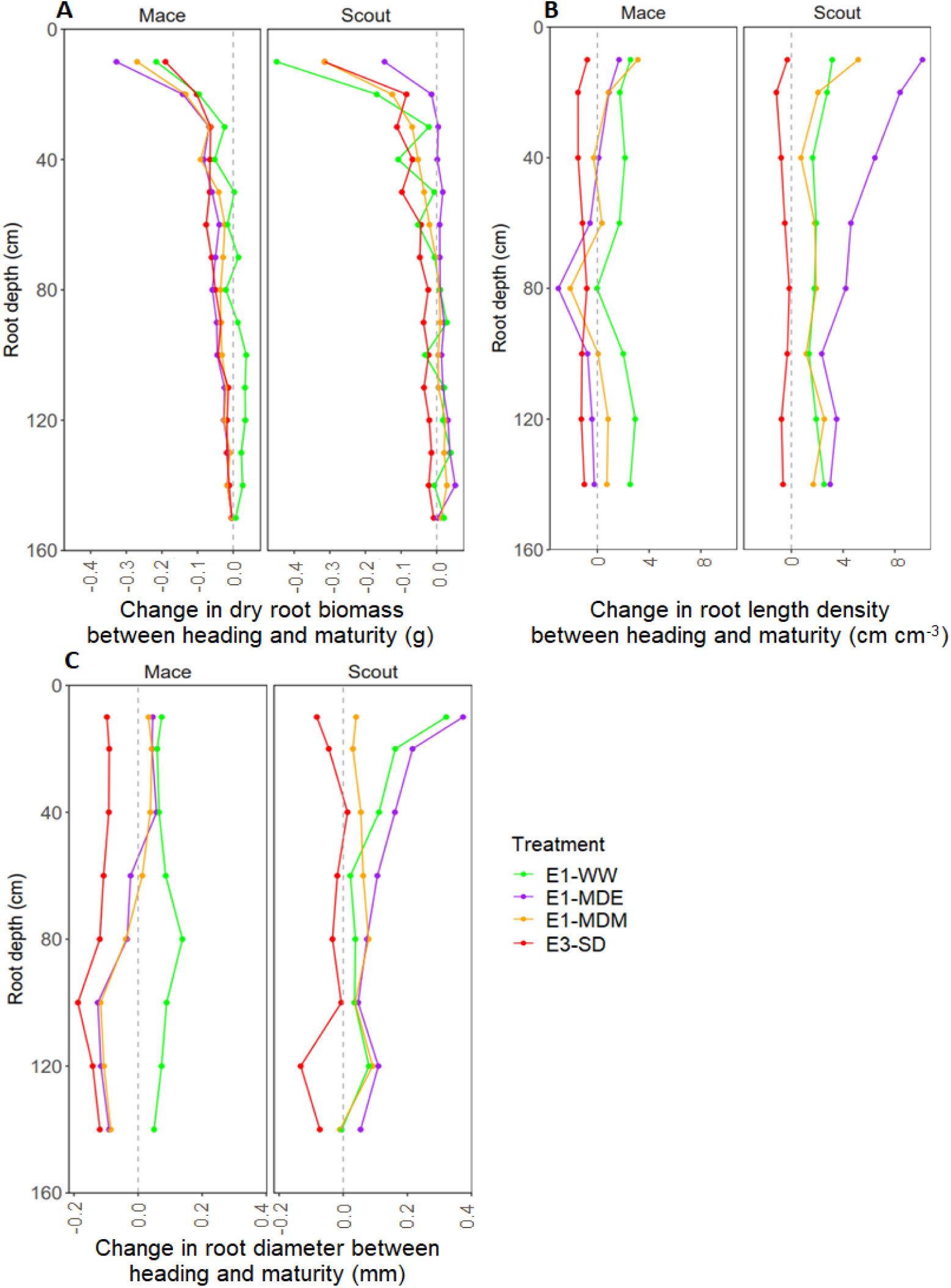
Change in root biomass (A), root length density (B), and root diameter (C) between heading (Z50) and maturity (Z92) for Mace and Scout at different soil depths (0 to 150 cm) in all studied water treatments. In each panel, the vertical dashed grey line corresponds to a value of zero representing no change between stages. Values to the left of the line (A) represent a decrease in biomass (net root senescence) while values to the right represent increase in biomass (net root growth).

Root length density in Scout was greater at maturity than at heading for all depths in all treatments except E3-SD where a small net senescence occurred (Fig. 7B). Roots of Scout also increased in diameter on average between heading and maturity for all treatments except E3-SD (Fig. 7C). For Mace, shallow and deep layer root length density between heading and maturity was reduced by water stress. Mace average root diameter was also reduced by water stress, except for shallow roots under moderate water stress which had a similar root diameter as under well-watered conditions (Fig. 7C, Supplementary Fig. 3C). The differences in root diameter between Scout and Mace were accentuated by all water stress treatments.

### 3.5 Water stress induced more canopy senescence in Mace than Scout

To address the question of whether root senescence observed for some water-stress treatments was related to canopy senescence, the greenness of the flag leaf was followed post anthesis using SPAD measurements (Fig. 8).

**Fig. 8.**
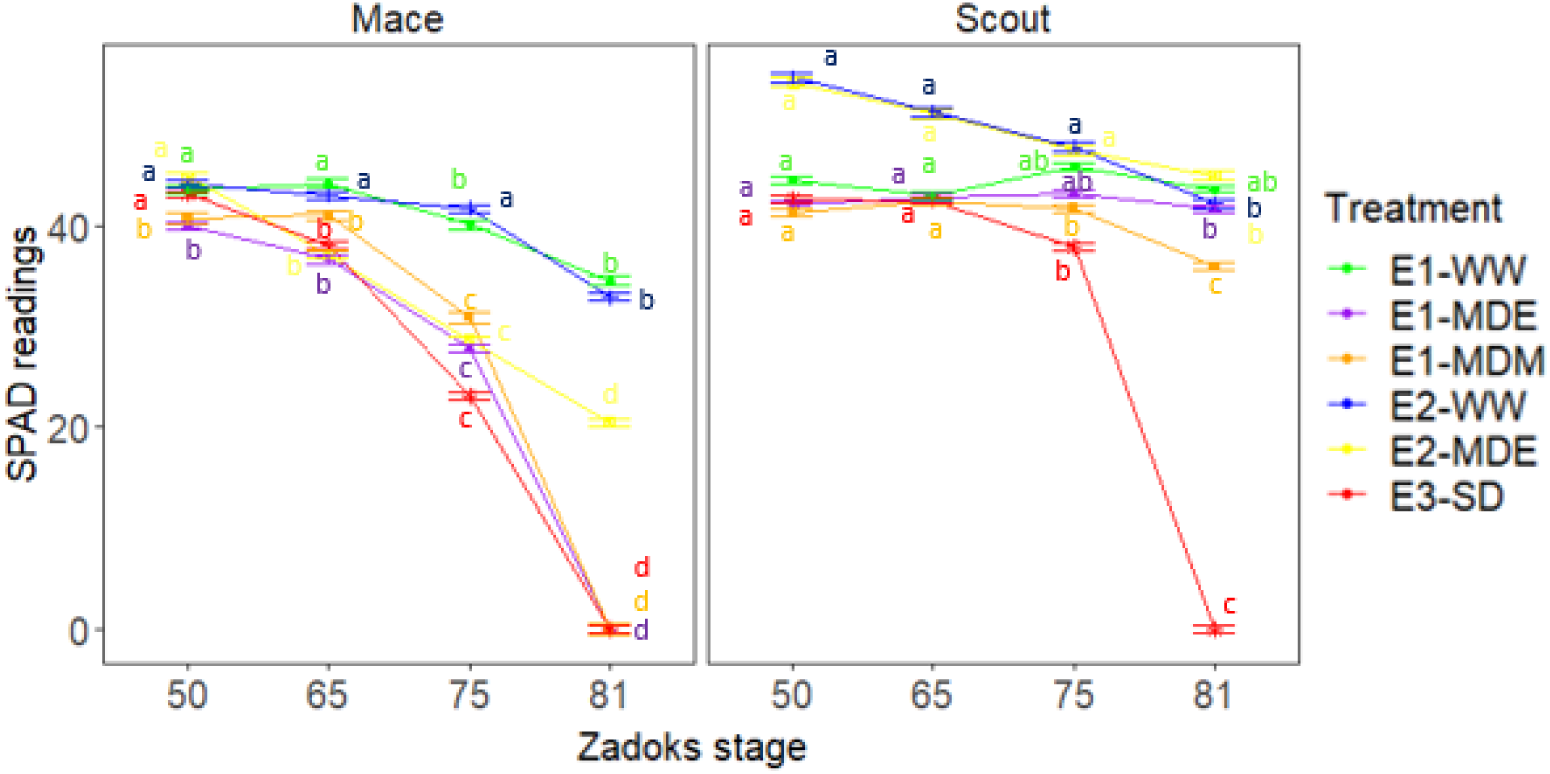
Dynamics of leaf greenness as indicated by SPAD values of Mace and Scout in all studied water treatments. SPAD values were measured in the centre of the flag leaf. Means that are significantly different (P<0.05) between Scout and Mace within each experiment are shown by the same letters above points. The analysis of variance was done separately for each experiment. Error bars represent the standard error of the mean (n=8).

Under well-watered conditions (E1-WW and E2-WW), both cultivars exhibited relatively high leaf SPAD values greater than 38 arbitrary SPAD units until late grain filling (Z81). In E1-WW, Scout had higher chlorophyl content than Mace at heading and anthesis and exhibited a small drop over the grain filling period, which was also observed in Mace (Fig. 8). Leaf area was also significantly higher for Scout than Mace at heading (Z50) under well-watered conditions (Table 1).

Moderate water stress (E1-MDE, E1-MDM, E2-MDE) resulted in accelerated leaf senescence during the grain filling (e.g., Z75) in Mace, while there was little or no effect on Scout flag leaf greenness (Fig. 8). Severe water stress (E3-SD) caused a rapid decrease in leaf greenness for both cultivars after anthesis (Z65) although Scout had significantly higher mean SPAD values than Mace up until mid-grain filling (Z75). Although not formally comparable, the mean leaf greenness (SPAD) for Scout at late grain filling (Z81) for E3-SD were reduced to a lower value than for any of the treatments in either experiment E1 or E2 for this genotype.

Mace showed leaf senescence from anthesis (Z65) to maturity (Fig. 8) which was correlated with loss of deep root biomass between heading (Z50) and maturity (Fig. 7) under moderate and severe stress. Similarly, leaf senescence in Scout correlated with root senescence in severe stress, while little or no senescence was observed in either deep roots or shoots of Scout under moderate stress (Figs. 7 and 8).

### 3.6 Water stress induced more yield loss in Mace than Scout

In well-watered conditions, the grain yield per plant in Mace (16 ± 1.5 and 12 ± 0.4 g in E1-WW and E2-WW, respectively) was similar to that of Scout (14 ± 1.5 and 13 ± 0.5 g; Table 1).

For moderate water stress, the yield per plant in Scout in treatments E1-MDE (16 ± 1.8 g), E1-MDM (12 ± 1.5 g) and E2-MDE (14 ± 0.4 g) remained similar to that in well-watered plants. In contrast, for Mace, the yield per plant was significantly lower for E1-MDE (7 ± 1.6 g), E2-MDE (8 ± 0.4), and E1-MDM (8 ± 1.8 g) compared with E1-WW and E2-WW, respectively (Table 1).

In severe water stress (E3-SD), yield of both genotypes was severely reduced compared with E1-WW or E2-WW with similarly low values of 2.5 ± 0.19 g for Mace and 3 ± 0.19 g for Scout (Table 1).

## 4. Discussion

The polyvinyl chloride (PVC) tube system was used to study the dynamics of root development at different depths under different water treatments using two wheat cultivars known to differ in seedling seminal root angle (Richard, 2017; Richard *et al*., 2018).

### 4.1 Moderate water stress induced root senescence or root growth depending on the genotype

In this study, Scout had similar or significantly greater shallow and deep root biomass than Mace at maturity (Figs. 2 and 5, Supplementary Fig. S1). Deep root biomass of Scout (i) significantly increased after anthesis under well-watered conditions (Fig. 2A, C), (ii) tended to slightly increase between heading and maturity when under moderate water stress (Fig. 5), but (iii) decreased post-heading in severe conditions (Figs. 3 and 7, Supplementary Fig. S4A). By contrast, shallow root biomass of Scout (i) senesced post-heading under well-watered, mid-grain filling moderate water-stress and severe water-stress conditions and (ii) tended to have a similar biomass at heading and maturity under early grain filling water stress, thus indicating some stress-induced root growth in these conditions. Overall, under water deficit conditions, Scout maintained deep roots more than the shallow roots (Fig. 7).

Conversely, Mace did not have post-anthesis deep root growth under well-watered conditions (Fig. 2), showed post-anthesis root senescence of shallow roots under well-watered conditions (Fig. 2), and showed shallow and deep root senescence for moderate and severe post-anthesis water stress treatments (Fig. 7). These observations suggest root development in Mace is more susceptible to water stress than Scout and may have adversely affected yield under the water-stress treatments examined. However, Mace is reputed to be well-adapted for the drought-prone Southern and Western Australian regions, where it has been widely grown (Ehdaie *et al*., 2012). Those regions have a Mediterranean climate characterized by winter-dominant rainfall (Singh *et al*., 2011) with medium to light soils, in which a quick finish to avoid terminal drought and heat may be more advantageous than developing post-anthesis root growth to extend the grain filling period. Such traits were observed in Mace in the current study.

### 4.2 Deep rooting as a drought adaptative trait

Earlier studies reported that Scout has a narrower seminal root angle phenotype than Mace (Alahmad *et al*., 2018; Richard *et al*., 2015). In other genotypes, this trait has been associated with development of deep roots and the ability to access water stored deep in the soil (Ludlow and Muchow, 1990). In agreement with this, in the current study, Scout also had a greater root biomass and root length density at depth than Mace, especially under moderate water stress (Fig. 5, Supplementary Fig. S1). These aspects of root architecture have been associated with drought tolerance, nutrient access, and yield through diverse mechanisms (Fukai and Cooper, 1995; Manschadi *et al*., 2006; Manske *et al*., 2000). Scout maintained root growth at depth under moderate water stress, while Mace showed net root senescence at depth in such conditions. As would be expected, severe prolonged water-stress treatment caused both root and leaf senescence for both genotypes. The effects of drought have been reported to be particularly detrimental when water stress extends through anthesis and grain-filling stages (Chenu *et al*., 2013; Farooq *et al*., 2014). Notably in the current study, water stress reduced the crop growth duration with increased senescence rate and an earlier maturity in Mace for moderate water stress and in both genotypes for severe water stress (Table 1). A shortened phenology is a well-known drought escape mechanism (Shavrukov *et al*., 2017). This strategy leads to a shortened crop cycle and is also characterised by decreased biomass due to the short time allowed for nutrient, light and CO_2_ uptake (Shavrukov *et al*., 2017), but can be suited for certain types of drought-prone environments (Collins and Chenu, 2021).

As can be expected, results indicate that adaptations favourable to moderate post-anthesis water stress in deep soils are not sufficient when the stress is too severe, as plants cannot survive without access to water. Deep rooting traits are also likely not as adaptive in soils with shallow and low water-holding capacity in regions where rainfall during the growing season is more important than deep stored moisture (Veyradier et al., 2013).

### 4.3 Maintenance of deep roots under moderate water stress was associated with a stay-green phenotype, a longer crop cycle and a greater plant yield for Scout

The greater deep root biomass observed for Scout at maturity, especially following moderate post-anthesis water stress, was associated with a stay-green phenotype, with plants retaining a higher green leaf area content until close to maturity (Fig. 8). In the severe water stress (E3-WD), Scout also retained leaf greenness later into development than Mace (Fig. 8). Such a stay-green phenotype with retention of both green leaf area and photosynthetic capacity for longer during grain filling, would be anticipated to maintain a higher yield as observed for different species in field conditions (Christopher *et al*., 2014; Nawaz *et al*., 2013; Thomas and Smart, 1993). Although it must be noted that photosynthetic capacity was not measured in the current study, the stay-green phenotype has been reported to enhance grain yield under post-anthesis drought stress in bread wheat (Adu *et al*., 2011; Bogard *et al*., 2011; Christopher *et al*., 2008; Lopes and Reynolds, 2012), durum wheat (Spano *et al*., 2003), and other cereals such as maize (Crafts-Brandner *et al*., 1984; Gentinetta *et al*., 1986; Zheng *et al*., 2009), rice (Ba Hoang and Kobata, 2009; Mondal *et al*., 1985; Wada and Wada, 1991), and sorghum (Borrell *et al*., 2023; Borrell *et al*., 2014). The ability to photosynthesise for extended periods, especially during the grain-filling stage may translate into increased above- and below-ground biomass accumulation leading to greater yield (Thomas and Howarth, 2000; Thomas and Smart, 1993) as observed for Scout in the current study (Table 1).

In the case of Scout, higher root biomass and root length density (e.g., Supplementary Figs. S1-S2) is likely to have resulted in more water uptake than Mace (although water uptake was not directly measured). If so, this helps to explain the higher SPAD readings, a proxy for chlorophyl content (Fig. 8) under the moderate post-anthesis water-stress treatments. This suggests that Scout adapted to seasonal drought stress through its root development and a delay in leaf senescence. Previous studies have highlighted that a greater ability to access deep water late in development can be associated with greater ability to extract water from deep in the soil late in the season when water use efficiency for grain production is high. This can have highly beneficial effect with a delay in senescence and enhanced productivity under terminal drought (Collins and Chenu, 2021; Kirkegaard *et al*., 2007; Manschadi *et al*., 2006).

Deep root growth, delayed root senescence and stay-green phenotype appear to have allowed Scout to maintain a longer period from anthesis to maturity compared to Mace in the moderately water-stressed (Table 1). Delay in progress to maturity, especially during the grain-filling stage is an important adaptive trait for cereal crops (Xiao *et al*., 2016) allowing longer time to accumulate biomass. In the present study, Scout reached maturity at a similar or slightly later time under moderate water stress than well-watered conditions while Mace matured earlier under the moderate water-stressed treatments leading to shortened anthesis to maturity duration (Table 1). Scout also had a greater yield per plant than Mace for these moderate stress treatments.

In the present study, Scout also had a greater root diameter than Mace both in well-watered and water-stressed conditions (Figs. 3, 6B and Supplementary Fig. S3). The diameter of deep roots in Mace decreased post-heading under moderate water stress (Fig. 6B) concomitantly with deep root biomass reduction (Fig. 5A). This suggests that as roots died, they become thinner. Decreased root diameter in wheat has previously been associated with water stress (Bakhshandeh *et al*., 2019).

Measuring root growth and viability in future could help understanding genetic variations for drought tolerance and assist breeding for better adapted genotypes.

## 5. Conclusion

The results of this study suggest that Scout maintained post-anthesis deep root growth under moderate post-anthesis water stress treatment whereas deep roots of Mace senesced. Deep root development potentially enabled access to water late in the season which likely aided Scout to maintain a stay-green phenotype and produce higher yield per plant under moderated water stress. Identification of such root traits that may enhance water uptake at depth late in crop development is anticipated to improve wheat adaptation to late season water stress conditions in drought-prone environments with heavy deep soils.

### Supplementary data

Supplementary Fig. S1. Dry root biomass at different soil depths in Mace and Scout for the different studied water treatments.

Supplementary Fig. S2. Root length density at different soil depths in Mace and Scout for the different studied water treatments.

Supplementary Fig. S3. Average root diameter at different soil depths in Mace and Scout for the different studied water treatment.

Supplementary Fig. S4. Change in dry root biomass, root length density and root diameter between heading and maturity at different depths in Mace and Scout for all studied water treatments.

## Acknowledgements

The authors are grateful for Renier Synman for technical help, and Bethany Rognoni and Michael Mumford for assistance in the statistical analysis.

## Author contributions

KS, JC and KC planned and designed the research. KS performed the experiments. KS and KC analysed the data. KS wrote the first version of the manuscript, with inputs from all the authors.

## Supplementary Figures

**Supplementary Fig. S3.**
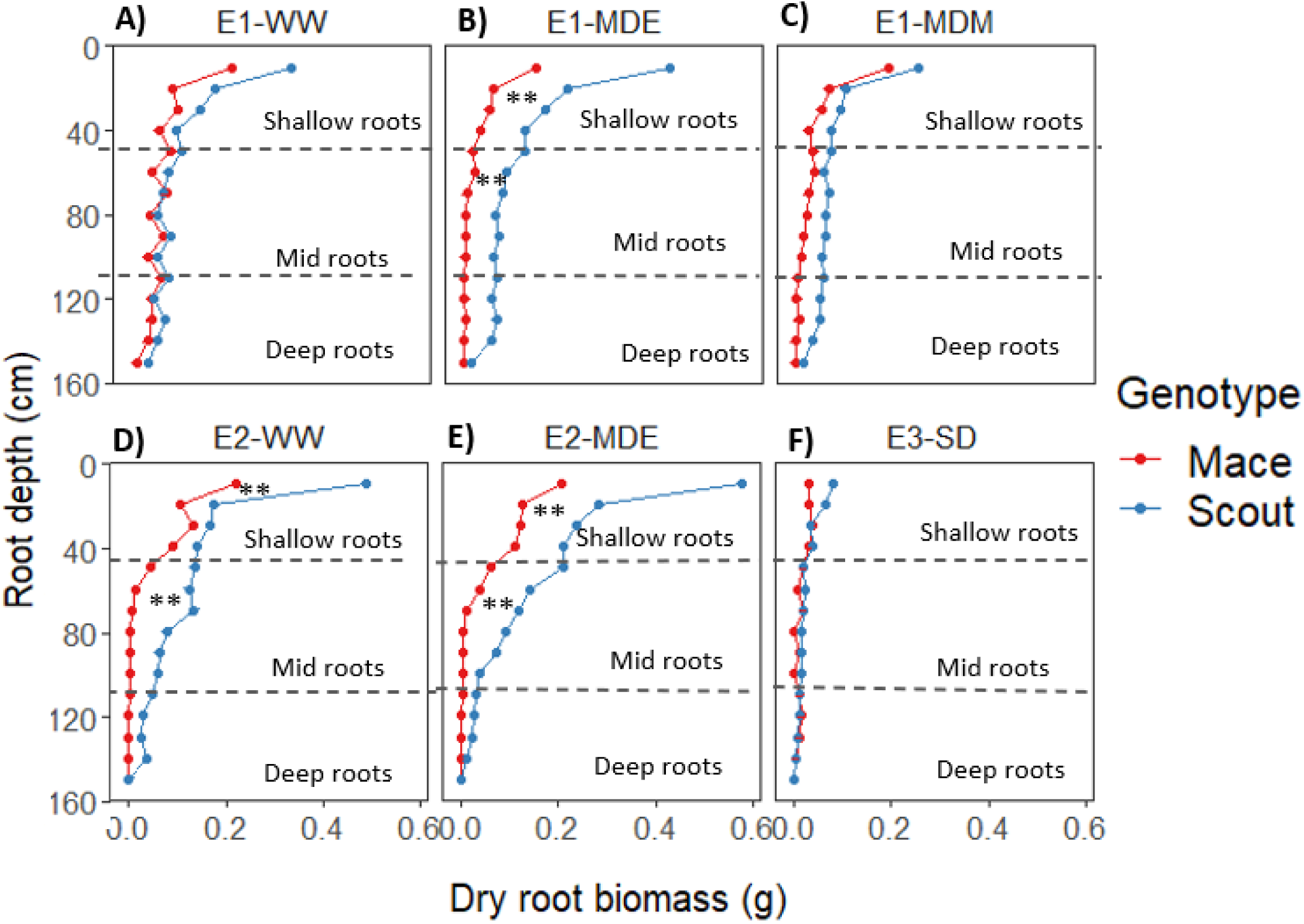
Dry root biomass at different soil depths (0 to 150 cm) in Mace and Scout for the different studied water treatments. Panels (A, B and C) correspond to water stress treatments in E1; (D, E) water-stress treatments in E2; and (F) sever water stress in E3. The analysis of variance was performed separately for each treatment. Asterisks indicate genotypic differences for shallow (0 to 50 cm), mid (50 to 100 cm) or deep (100 to 150 cm) roots (P<0.01). This figure presents data from Figure 6A treatment by treatment.

**Supplementary Fig. S4.**
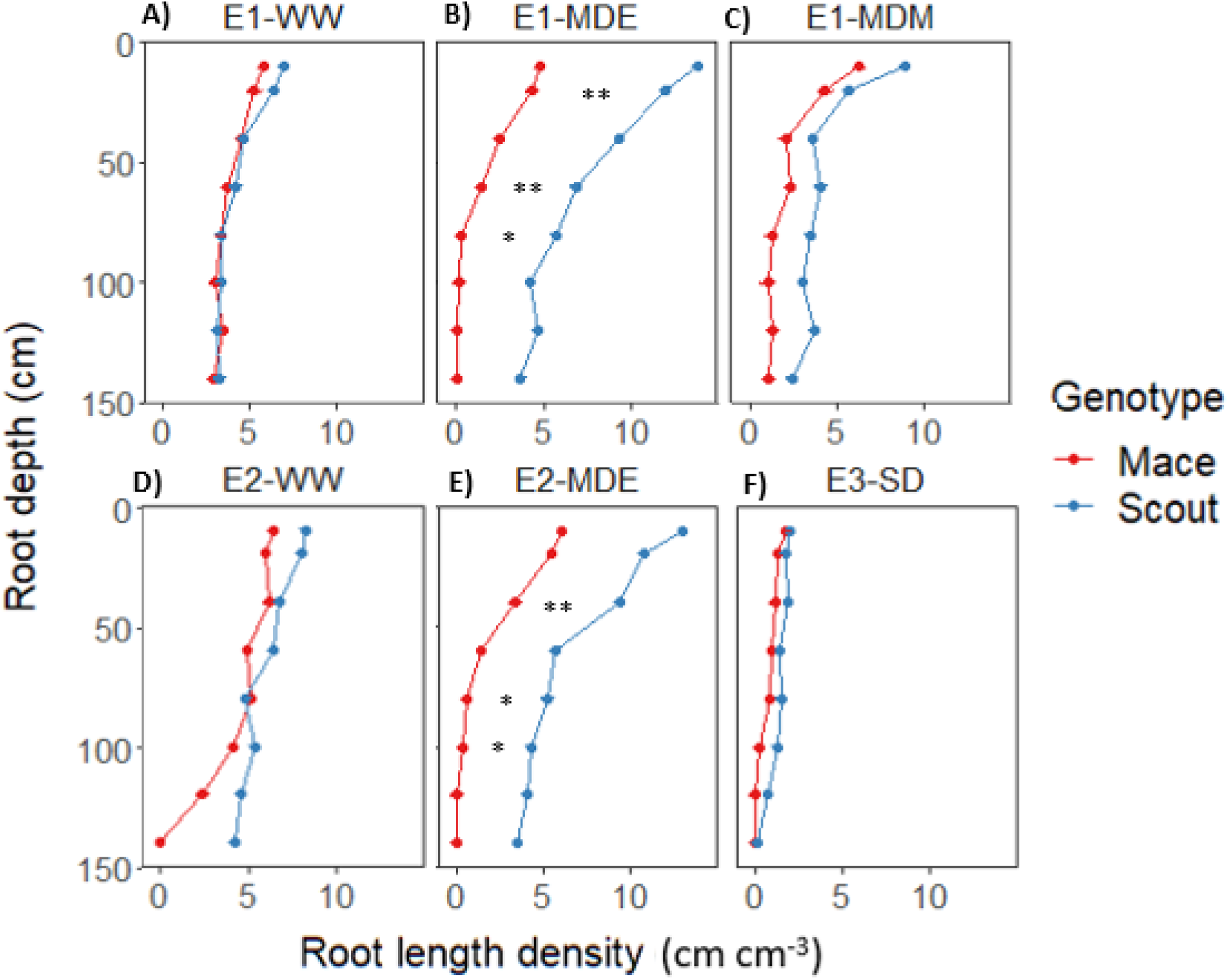
Root length density at different soil depths in Mace and Scout for the different studied water treatments. Panels (A, B and C) correspond to water stress treatments in E1; (D, E) water-stress treatments in E2; and (F) sever water stress in E3. The analysis of variance was performed separately for each treatment. Asterisks indicate genotypic differences for roots at a specific depth (P<0.01). This figure presents data from Figure 7A treatment by treatment.

**Supplementary Fig. S5.**
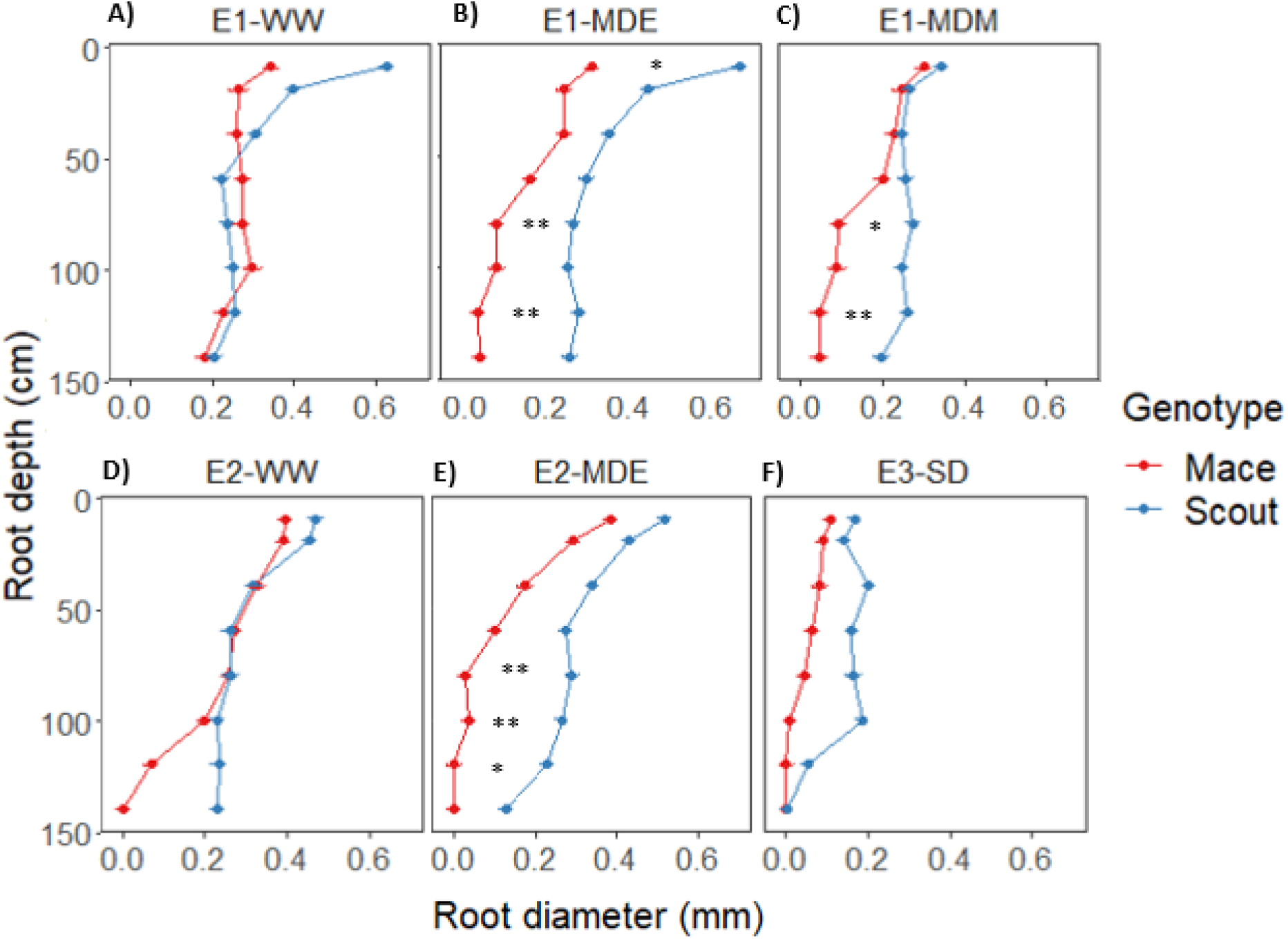
Average root diameter at different soil depths in Mace and Scout for the different studied water treatment. Panels (A, B and C) correspond to water stress treatments in E1; (D, E) water-stress treatments in E2; and (F) sever water stress in E3. The Analysis of variance was performed separately for each treatment. Asterisks indicate genotypic differences for roots at a specific depth (P<0.01). This figure presents data from Figure 7B treatment by treatment.

**Supplementary Fig. S6.**
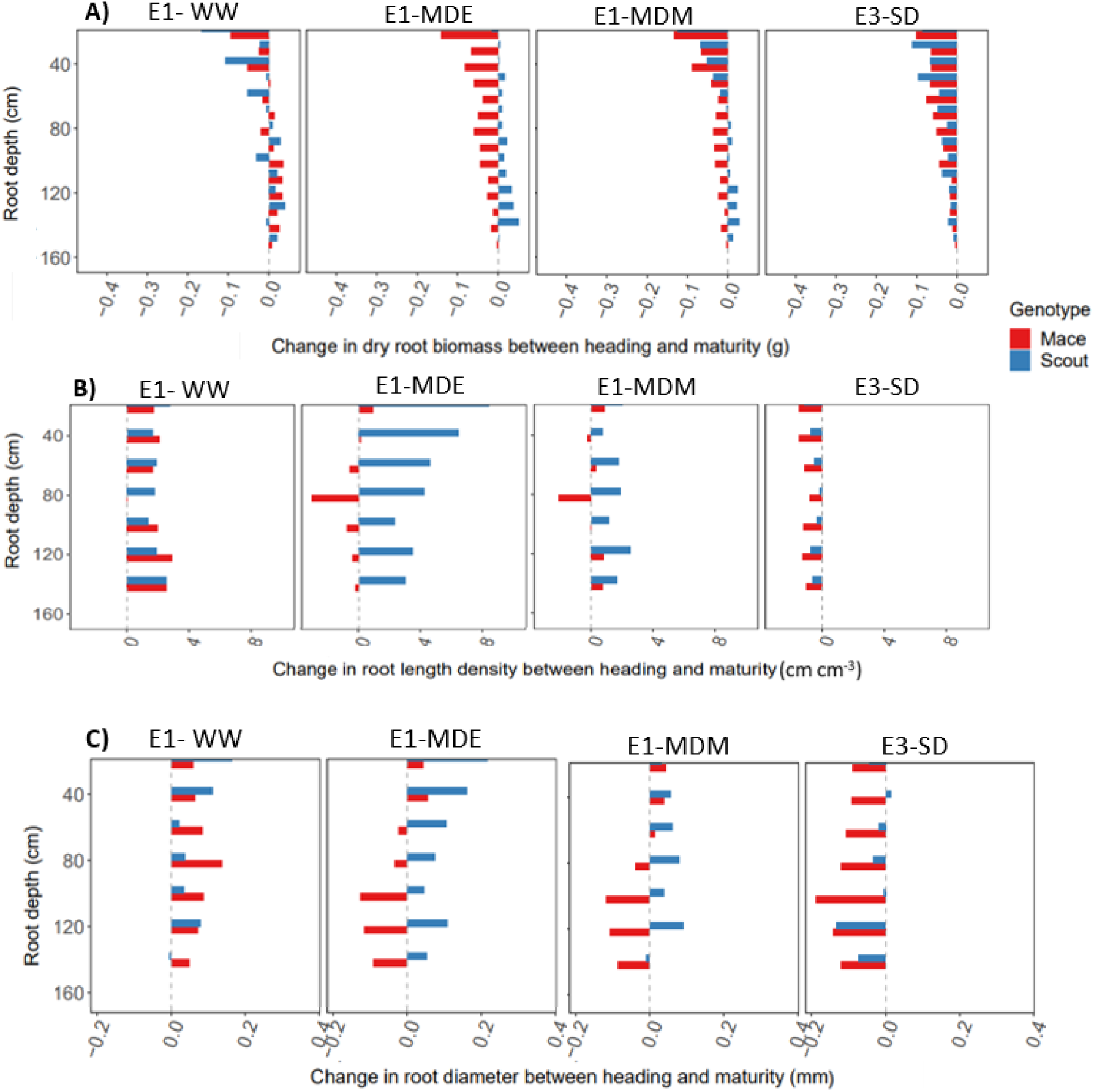
Change in (A) dry root biomass, (B) root length density and (C) root diameter between heading and maturity at different depths (0 to 150 cm) in Mace and Scout for all studied water treatments. This figure presents data from Figure 8 treatment by treatment.

